# Laboratory evolution from social to solitary behavior in the N2 reference strain is unnecessary for its fitness advantages

**DOI:** 10.1101/309997

**Authors:** Yuehui Zhao, Lijiang Long, Wen Xu, Richard F. Campbell, Edward L. Large, Joshua S. Greene, Patrick T. McGrath

## Abstract

The standard reference *Caenorhabditis elegans* strain, N2, has evolved marked behavioral changes since its isolation from the wild 67 years ago. Laboratory-derived variation in two genes, *npr-1* and *glb-5*, suppress aerotaxis behaviors on food, resulting in N2 animals evolving from social to solitary feeding strategies. We show here that the derived alleles of *npr-1* and *glb-5* can confer large fitness advantages in standard laboratory conditions, suggesting that the changes in feeding strategies were beneficial to the N2 strain. However, by using environmental manipulations that suppress social behaviors, we showed the fitness advantages of the derived alleles remained unchanged, suggesting selection on these alleles acted through biological traits unrelated to solitary behavior. Transcriptomics analysis, developmental timing assays, and feeding assays showed that N2 animals mature faster, produce more sperm, and eat more food than a strain containing ancestral alleles of these genes (CX12311) regardless of the behavioral strategies. The O2-sensing neurons URX, AQR, and PQR and the pheromone biosynthesis and lipid regulating enzyme encoded by *daf-22* are necessary for the full fitness advantages. We suggest that changes to social/solitary behavior in N2 were a pleiotropic consequence of *npr-1* and *glb-5’s* ability to modify integrated O2 and pheromone neural circuits that regulate feeding rate and reproductive development. Together, our results demonstrate how laboratory evolution can lead to profound changes in a strain used as a model by for understanding a variety of fundamental biological processes.

## Introduction

Since the fundamental work of Gregor Mendel elucidating the laws of genetic transmission, model organisms have enabled experimenters to gain fundamental insights into many biological processes. Modern research tools are facilitating the use of new and unusual species to analyze longstanding biological questions (Alfred and Baldwin 2015; Gladfelter 2015; Goldstein and King 2016; Russell et al. 2017). More and more species are reared in the laboratory as models for biological traits of interest. An issue for these approaches, particularly for comparative analysis or for those addressing evolutionary questions, is the extreme shift in environment, and associated selective pressures, these populations experience. All species evolve through the process of natural selection and genetic drift; many model organisms have evolved by exposure to the novel and artificial conditions experienced in the lab (Orozco-terWengel et al. 2012; Duveau and Felix 2012; Goto et al. 2013; Kasahara et al. 2010; Marks et al. 2010; Stanley and Kulathinal 2016; Yvert et al. 2003). Understanding the process of adaptation of wild populations to captivity is necessary to understand how the genetic, developmental, and neural circuits are changed in these laboratory populations. Additionally, these types of studies can be useful for understanding basic evolutionary processes (Fisher and Lang 2016; Lenski 2017; Teotonio et al. 2017). For example, the ability to manipulate model organisms in the lab provides a greater opportunity to test adaptive hypothesis beyond arguments of plausibility and address the role of pleiotropy and other competing themes in the evolution of biological traits (Gould and Lewontin 1979).

As a model for understanding laboratory adaptation in a multicellular organism, we have focused our studies on the N2 strain of *Caenorhabditis elegans*. N2 is the canonical reference strain used by thousands of *C. elegans* labs across the world. While this strain was introduced to the genetics research community by Sydney Brenner in 1974 (Brenner 1974), it was actually isolated by L.N. Staniland and Warwick Nicholas from mushroom compost in 1951, spending multiple decades (~300-2000 generations) in two primary growth conditions: on agar plates where bacteria was its primary food source or in liquid axenic media (Sterken et al. 2015). A small number (~100) of new mutations that arose and fixed in the N2 strain following isolation from the wild have been identified (McGrath et al. 2011), including a neomorphic, missense mutation in the neuropeptide receptor gene *npr-1* and a recessive, 765 bp duplication in the nematode-specific globin gene *glb-5*. These mutations were originally identified for their role in foraging and aerotaxis behaviors and were initially thought to represent natural genetic variants (de Bono and Bargmann 1998; Persson et al. 2009). A large body of work has found that these genes regulate the activity of the URX-RMG neuronal circuit that controls O_2_ responses on food (Chang et al. 2006; Coates and de Bono 2002; Gray et al. 2004; Macosko et al. 2009; McGrath et al. 2009; Persson et al. 2009). Animals with the ancestral alleles of *npr-1* and *glb-5* follow O_2_ gradients to the border of bacterial lawns and feed in groups (called social behavior); animals containing the derived alleles of these genes ignore O_2_ gradients in the presence of food and feed alone (called solitary behavior) (**Figure 1a**) (Gray et al. 2004).

**Figure 1.**
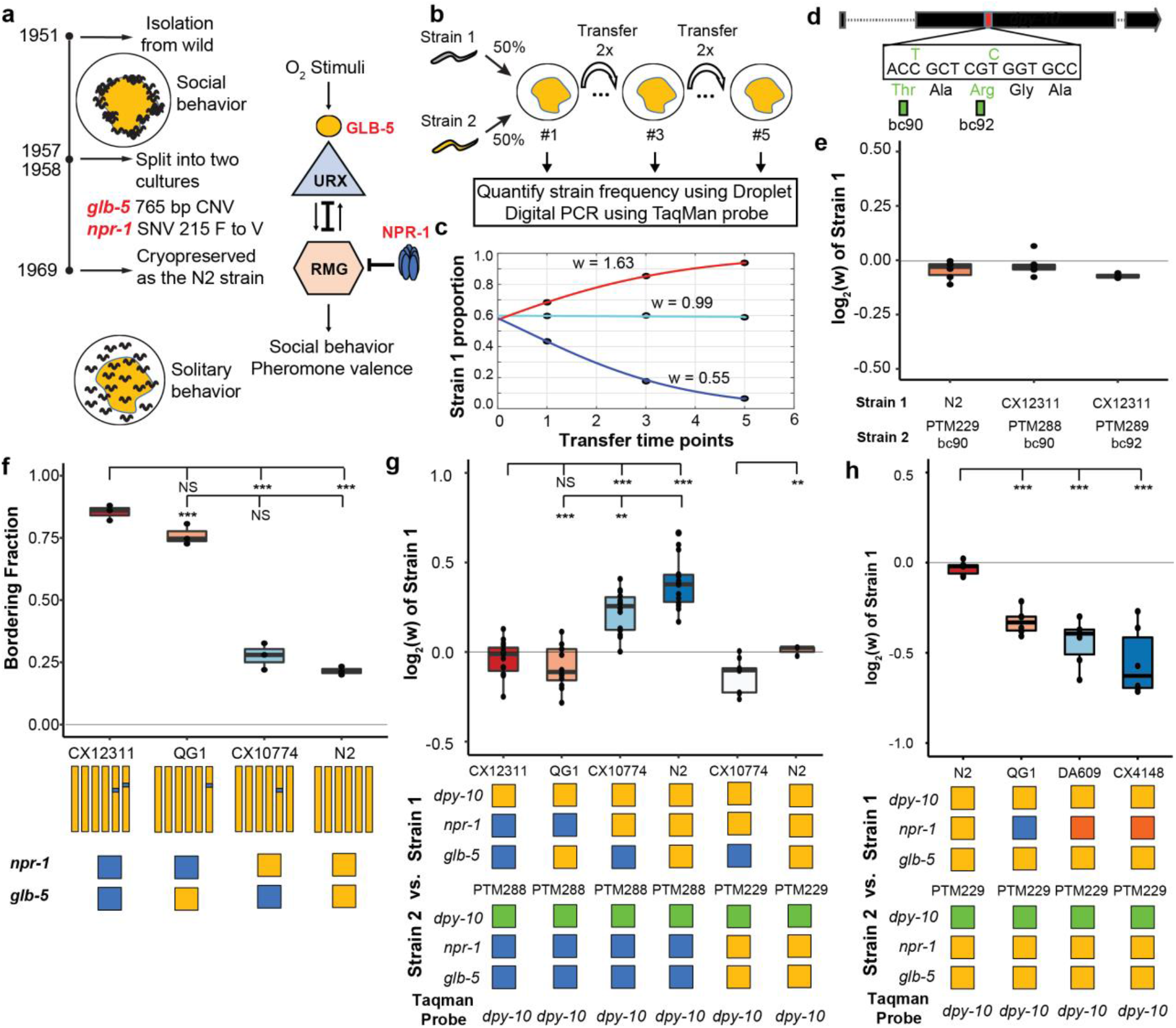
Selective advantage of derived alleles of *npr-1* and *glb-5*. **a**. Derived alleles in *npr-1* and *glb-5* arose after isolation from the wild and caused changes in foraging behavior in the N2 reference strain. These genes regulate the activity of the URX/RMG neural circuit. **b**. Schematic of pairwise competition experiments used throughout the paper to quantify fitness differences between two strains. **c**. Relative proportion of each strain as ascertained by Droplet Digital PCR (dots) is used to estimate relative fitness (line). **d**. Silent mutations were edited into the 90^th^ or 92^nd^ amino acid of the *dpy-10* gene using CRISPR/Cas9 to create a common SNV for Droplet Digital PCR. We refer to these as barcoded strains. **e**. Competition experiments between the parent strain (top) and the same strain containing one of the silent mutations. We display each competition experiment as a dot overlaid on top of a boxplot showing the mean, first, and third quartiles. **f**. The bordering rate of the N2 reference strain compared to three NIL strains containing ancestral alleles of *npr-1* and/or *glb-5* introgressed from the CB4856 wild strain. Schematic of each NIL shown below along with the allele of *npr-1* and *glb-5* they contain. Orange represents N2-derived DNA and blue represents CB4856-derived DNA. To ascertain statistical significance, ANOVA was used followed by a Bonferonni correction for multiple tests. **g**. Competition experiments between NILs shown in panel f against barcoded strains shown in panel **e**. Green box indicates the strain contains the barcoded allele of *dpy-10*. Positive values indicate Strain 1 is more fit; negative values indicate Strain 2 is more fit. **h**. Competition experiments between strains containing two loss-of-function alleles of *npr-1* (red boxes) along with controls.

We have previously proposed that the derived alleles of *glb-5* and *npr-1* were fixed by selection as solitary animals are more likely to be picked when propagating animals to new plates (McGrath et al. 2009). However, aggregation behavior in the ancestral *npr-1* strain appears to create local food depletion leading to a weak starvation state, which reduces reproduction and growth (Andersen et al. 2014). Potentially this starvation difference could be responsible for the fitness differences of the strains. Consistent with both hypothesis, a number of experimental crosses or competition experiments between parental strains that are polymorphic for *npr-1* have resulted in enrichment of the derived allele of *npr-1*, suggesting it confers a fitness advantage under standard lab husbandry (Gloria-Soria and Azevedo 2008; Noble et al. 2017; Weber et al. 2010).

In order to distinguish between these hypothesis, we performed pairwise competition experiments following a number of environmental and/or genetic manipulations. Surprisingly, our results suggest that neither hypothesis is correct. While the derived *npr-1* and *glb-5* alleles increase fitness of animals on agar plates, the differences in social vs. solitary behavior are not necessary for their differences in fitness. Instead, our work suggests that fitness gains are due to increases in feeding rates, mediated by changes to the neural circuits that control pheromone responses. Our work demonstrates that even when alleles are identified that confer fitness advantages, care must be taken in inferring the phenotypes that are responsible due to the pleiotropic actions of genetic changes.

## Results

### Derived alleles of *npr-1* and *glb-5* increase fitness in laboratory conditions

In previous reports, we have used multigenerational pairwise competition experiments to compare the relative fitness of two strains (**Figure 1b**) utilizing Droplet Digital PCR with a custom TaqMan probe to quantify the proportion of each genotype (Evans et al. 2017; Greene et al. 2016; Large et al. 2016). To quantify this assay, we used a generic selection model to estimate the relative fitness difference (w) between the two strains (**Figure 1c**). In this context, relative fitness measures the generational change in relative abundance of each of the two strains. We also used CRISPR-enabled genome engineering to create strains with a silent mutation in the *dpy-10* gene using a previously published high-efficiency guide RNA (**Figure 1d**) (Arribere et al. 2014), which we will refer to as barcoded strains. These barcoded strains allow us to use a common Taqman probe to quantify the relative fitness of a test strain against these barcoded strains. We confirmed that the *dpy-10* silent mutation had no statistically significant effect on fitness in two genetic backgrounds studied throughout this report (**Figure 1e**).

In order to test the fitness effect of the derived alleles of *npr-1* and *glb-5*, we utilized three previously described near isogenic lines (NILs) containing ancestral alleles of *npr-1* (QG1), *glb-5* (CX10774), or both genes (CX12311) introgressed from the CB4856 wild strain into the standard N2 background (Bernstein and Rockman 2016; McGrath et al. 2009; McGrath et al. 2011). The *npr-1* introgressed region is ~110 kb in size and the *glb-5* introgressed region is ~290 kb in size. For brevity, we will refer to genotype of these introgressed regions throughout this report by the ancestral/derived allele they contain (e.g. the ancestral allele of *npr-1* vs the introgressed region containing the ancestral allele of *npr-1)*. In contrast to the N2 strain, the CX12311 strain aggregates at the border of bacterial lawns where O_2_ levels are lowest due to the increased height of the bacterial lawn. We confirmed previous reports that both the derived alleles of *npr-1* and *glb-5* suppress bordering behavior to varying degrees (Bendesky et al. 2012; de Bono and Bargmann 1998; McGrath et al. 2009); *npr-1* accounted for the majority of the difference with *glb-5* playing a modulatory role (**Figure 1f**). To compare the relative fitness of the four strains, we competed each strain against the barcoded CX12311 strain, transferring animals each generation by washing to minimize potential sources of investigator bias towards picking social or solitary animals (**Figure 1g**). The N2 strain was the most fit in these conditions, with a relative fitness (w) of ~1.30. Interestingly, the fitness effects of the *glb-5* and *npr-1* regions were epistatic - the derived allele of *glb-5* increased the relative fitness in the derived *npr-1* background but showed no effect in the ancestral allele of *npr-1*. The derived *npr-1* allele increased fitness in both backgrounds of *glb-5*. To confirm the fitness advantage of the derived *glb-5* allele in the derived *npr-1* background, we also competed the CX10774 NIL against the barcoded N2 strain (**Figure 1g**). The estimated selective coefficient (a common measure of the fitness difference of a beneficial allele) of the *glb-5* allele in the *npr-1* derived background was s = 0.10 (0.06 – 0.13 95% confidence interval), the estimated selective coefficient of the *npr-1* allele in the *glb-5* ancestral background was s = 0.17 (0.12 - 0.23 95% confidence interval), and the estimated selective coefficient of the npr-1 allele in the *glb-5* derived background was s = 0.30 (0.27 - 0.34 95% confidence interval). These selective coefficients are comparable to beneficial alleles identified in other organisms, such as the haplotype responsible for lactase persistence (~0.01-0.19) (Bersaglieri et al. 2004) and the sickle-cell trait (0.05 – 0.18) in humans (Li 1975).

While the introgressions surrounding the *npr-1* and *glb-5* genes are relatively small, these NIL strains carry additional polymorphisms in surrounding genes from the CB4856 strain. We also performed competition experiments using two previously published *npr-1* loss-of-function alleles *(ad609* and *ky13)* (de Bono and Bargmann 1998) against the N2 barcoded strains. Both the *npr-1(ad609)* and *npr-1(ky13)* loss-of-function alleles decreased the animal’s relative fitness in an amount comparable to the ancestral allele (**Figure 1h**). We did not perform similar experiments on the *glb-5* gene. Altogether, our work suggests that the *npr-1* derived allele increases fitness of animals in laboratory conditions and also suggests that the derived allele of *glb-5* increases the fitness of animals in a *npr-1* dependent manner.

### Suppression of foraging behavior differences between N2 and CX12311 does not suppress their fitness differences

Animals with reduced function of *npr-1* sense environmental O_2_ levels and aerotax towards their preferred O_2_ levels (10%) in the presence of foods which results in aggregation of animals at the borders of the lawn (Chang et al. 2006; Cheung et al. 2005; Gray et al. 2004). This behavior can be suppressed by lowering environmental O_2_ levels to the animals preferred O_2_ concentrations (Gray et al. 2004). We decided to use this environmental manipulation to test the hypothesis that the social foraging behavior was necessary for the fitness disadvantage experienced by strains containing the ancestral alleles of *npr-1* and *glb-5*. Our above experiments hinted that this hypothesis might be incorrect as the derived *glb-5* allele reduced bordering behavior in the ancestral *npr-1* background without an associated increase in fitness. We first confirmed that we could suppress the bordering behavior differences between CX12311 and N2 by reducing environmental O_2_ levels to 10% or 3% using a Biospherix chamber (**Figure 2a** and **Videos S1-4**). CX12311 animals did not form any social groups in the center of the lawn at the lowered O_2_ levels and were indistinguishable from N2 by visual inspection. Despite the behavioral similarity of these animals at these lower O_2_ levels, the relative fitness differences between the N2 and CX12311 strains remained (**Figure 2b**). To further confirm that aggregation behavior was not necessary for the fitness differences, we also performed competition experiments on uniform bacterial lawns (UBLs), which are constructed so that the entire plate is covered with a thin bacterial lawn to remove the O_2_ gradients created by the unequal thickness of bacteria in normal lawns. UBLs have been used to suppress *npr-1*-dependent differences in survival in response to bacterial pathogens (Reddy et al. 2009), however, the UBLs were unable to suppress the fitness advantage of N2 animals (**Figure 2c**).

**Figure 2.**
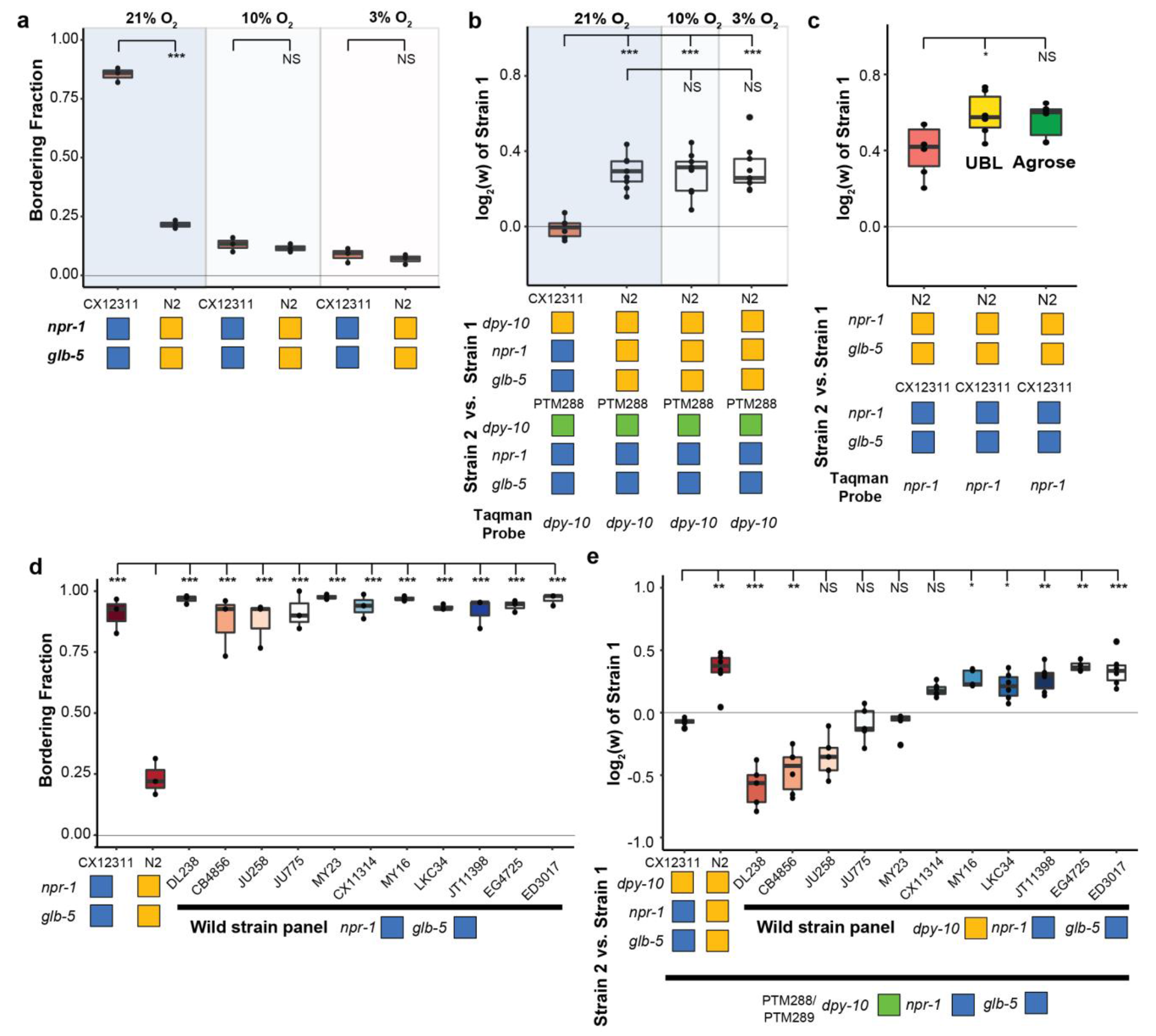
Fitness advantage of N2 is independent of foraging behavior. **a**. Environmental O_2_ levels were manipulated using a Biospherix chamber. Bordering behavior was suppressed in CX12311 at 10% or 3% environmental O_2_ levels. **b**. Fitness advantages of N2 over the barcoded CX12311 strain were independent of environmental O_2_. **c**. Fitness advantages of N2 were also present on uniform bacterial lawns (UBL) where animals were unable to border and on plates containing agarose which prevent burrowing behaviors. **d**. A panel of 11 wild strains was tested for bordering behavior. Each of these wild strains contains ancestral alleles of *glb-5* and *npr-1*. **e**. Competition experiments between 11 wild strains and barcoded CX12311 animals. Despite the similarity of bordering behavior, these wild strains displayed a range of relative fitness.

Animals that carry the ancestral *npr-1* allele can burrow into agar when food is depleted (de Bono and Bargmann 1998), raising the possibility that the fitness gains of N2 could be a result of the transfer process, which selects for animals on the surface of plates. While visual inspection of the two strains at 10% and 3% did not reveal any obvious differences in burrowing behavior, we also tested the role of burrowing in the fitness differences more rigorously by using modified nematode growth plates that contain agarose that prevents burrowing (Andersen et al. 2014). The relative fitness differences between N2 and CX12311 remained unchanged (**Figure 2c**).

These experiments motivated us to also test the relative fitness differences of eleven other wild strains isolated from different parts of the world using strains provided by the *C. elegans* Natural Diversity Resource (Cook et al. 2017). Each strain was competed against a barcoded CX12311 strain. Consistent with their *npr-1* genotype, these wild strains all aggregated at the borders of the bacterial lawn (**Figure 2d**), but their relative fitness differences varied wildly (**Figure 2e**). The relative fitness of two of the strains (CB4856 and DL238) was greatly reduced compared to the CX12311 strain. The relative fitness of five of the strains were comparable to the N2. The relative fitness of the remaining four strains was statistically indistinguishable from CX12311. These results further support our previous results that social behavior is not the major determinant of fitness levels in laboratory conditions.

### Development speed and spermatogenesis are increased in N2 in an O_2_-independent manner

To gain more insight into the phenotypes that could be responsible for the fitness increases of the N2 strain, we performed RNA sequencing to analyze the transcriptomes of bleach-synchronized N2 and CX12311 animals grown in either 10% O_2_ or 21% ambient O_2_ levels. Animals were allowed to develop to the L4 stage and harvested at identical times. We first performed Principal Component Analysis (PCA) analysis to analyze how the environmental and genetic differences globally regulated the transcriptomes of the animals. If environmental O_2_ and the genetic background had independent effects on the transcriptomes, we expected to find two major components in the PCA analysis. However, the PCA analysis identified a single component that explained the majority of the variance (77.9%). The genetic and environmental perturbations had similar additive effects on the first component in an additive manner (**Figure 3a**). Reducing O_2_ levels from 21% to 10% had the same effect on the transcription profiles as changing the background from CX12311 to N2. Consequently, the animals that differed in both genetic background and environmental O_2_ levels (N2 – 21% O_2_ vs CX12311 – 10% O_2_) also showed the most similar transcriptional profiles. These patterns were also seen in Hierarchical Clustering using the 1202 differentially-expressed genes (**Figure 3b**). These results suggest that the foraging behavioral differences are not responsible for the underlying transcriptomics differences between the different strains and environmental conditions.

**Figure 3.**
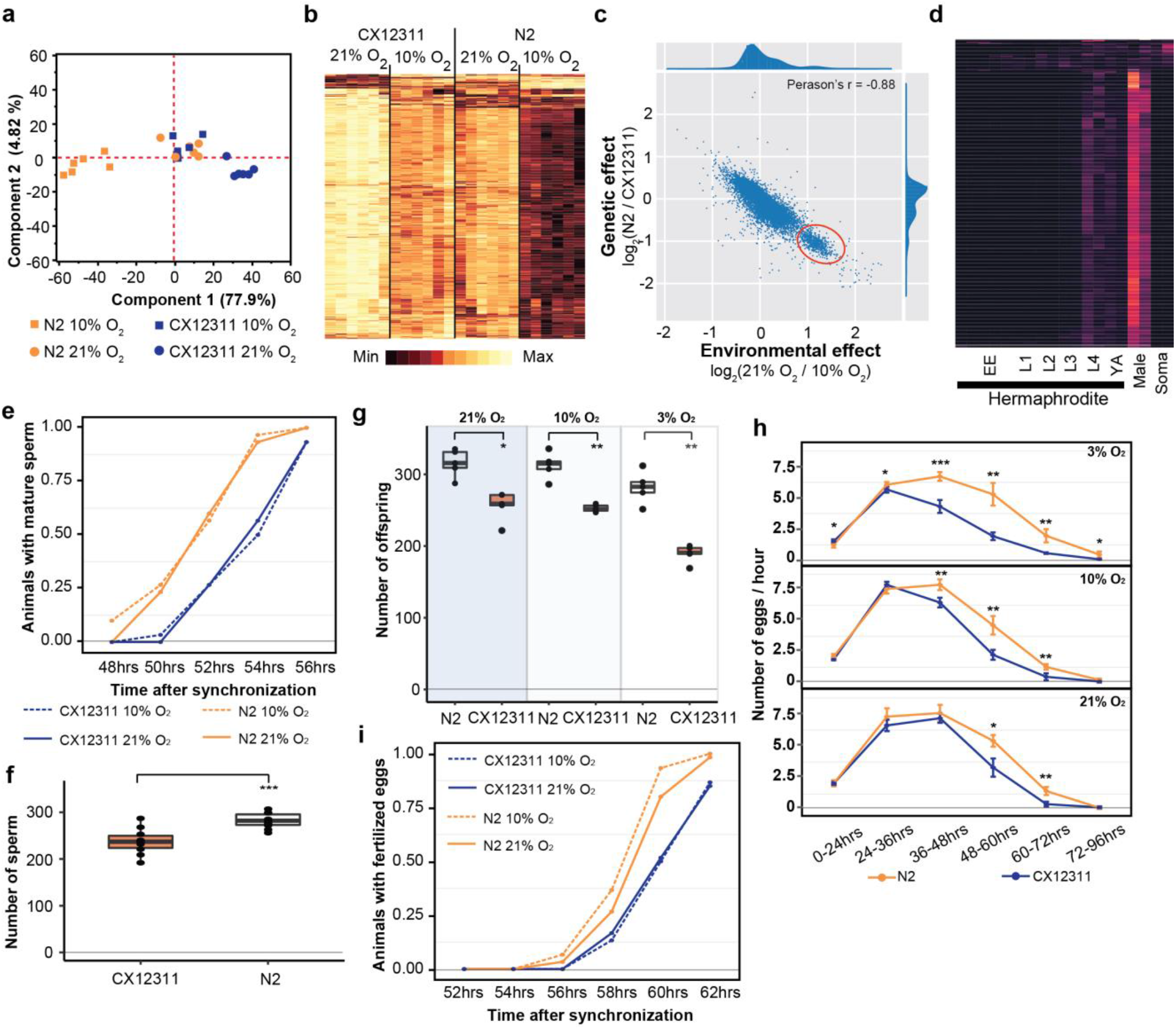
Reproductive timing in N2 occurs earlier than the CX12311 strain. **a**. PCA analysis of transcriptional profiles of bleach-synchronized N2 and CX12311 animals grown in 10% or 21% environmental O_2_ (six replicates per strain/condition). The largest two eigenvectors are shown, along with the amount of variance they explain. Developmental age of animals is approximately L4 stage. **b**. Hierarchical clustering of normalized, differentially expressed genes. Columns show strain and conditions; rows show gene expression. **c**. Averaged effect of genotype (y-axis) vs environment (x-axis) for each differentially expressed gene. A small cluster of 652 genes with similar changes is circled in red. **d**. The developmental expression of these 652 genes was further investigated using a previously published dataset. Columns show developmental stage and rows show each gene. Most of these gene peaked in expression in L4 hermaphrodite animals and was further enriched in male L4 animals (Male). Soma indicates expression levels from somatic cells, suggesting this cluster is enriched in germline cells. **e**. Animals identified with mature sperm. x-axis indicates time since synchronization using hatch-off. Strain/condition shown in legend. **f**. Number of sperm produced by each strain as determined by DAPI straining. **g**. Averaged total number of offspring produced by each strain when grown in different environmental O_2_ levels. **h**. Averaged egg-laying rate of L4-synchronized N2 and CX12311 animals when grown at different O_2_ levels. x-axis indicates time since L4 stage. **i**. Number of animals observed with fertilized eggs in their uterus. x-axis indicates time from synchronized egg-lay.

The effects of the derived *npr-1* and *glb-5* alleles mimics the effects of lowering environmental O_2_ from 21% to 10%. To further gain insight into this connection, we plotted the average transcriptional change between the strain backgrounds vs the average transcriptional change between the environmental O_2_ concentrations for each gene (**Figure 3c**). Surprisingly, we observed a bimodal distribution of values, with a cluster of 652 genes centered at 1.2 log_2_-fold change (**Figure 3c** – red circle). This is unexpected, as it suggests that the environmental and genetic perturbations had identical effects on transcription for all of these genes. When we inspected this list of genes, we noticed a large number of genes that are known to be involved in spermatogenesis. We further investigated the developmental regulation of these 652 genes using previously published transcriptomics data isolated from hermaphrodites or males at specific developmental time points (Boeck et al. 2016) (**Figure 3d**). The expression of the majority of these genes peaked during the L4 stage in hermaphrodites, was further enriched in L4 males, and suppressed in somatic cells isolated from L4 animals. These observations are consistent with this cluster of genes being involved in spermatogenesis, which occurs during the L4 stage (when RNA was isolated) in hermaphrodite animals.

We reasoned that the transcriptomics data could indicate a difference in the relative timing of spermatogenesis and/or the number of sperm that are produced in each genetic background/environmental condition. L1 larval stage animals were synchronized; subsequent differences in developmental speed would result in animals in slightly different stages of L4. To test this, we synchronized CX12311 and N2 animals, placed them in 10% or 21% environmental O_2_, and identified the number animals containing mature sperm at 2-hour intervals from 48-56 hours. N2 animals began spermatogenesis approximately two hours earlier than the CX12311 animals, regardless of the environmental O_2_ levels (**Figure 3e**). Hermaphrodites undergo spermatogenesis for a fixed period of time before permanently switching gametogenesis to the production of oocytes, resulting in the development of a fixed number of self-sperm that are stored in the spermathecal (Hubbard and Greenstein 2005). To test whether these strains produced the same number of sperm, we used DAPI staining to count the number of sperm found in the spermathecal. Not only did N2 animals start spermatogenesis earlier, they also produced more sperm (**Figure 3f**). The total fecundity of N2 hermaphrodites that do not mate with males is determined by the number of self-sperm. We confirmed that the difference in self-sperm number also resulted in a larger overall brood size (**Figure 3g**) and as expected from computational modeling (Large et al. 2017), an increased rate of egg-laying later on in life (**Figure 3h**).

The timing of sexual maturity is an important factor in determining the fitness of animals. We also tested whether the differences in timing of spermatogenesis could lead to differences in when fertilized eggs are produced. We performed similar experiments as above and monitored the time fertilized eggs could be observed in the uterus at two-hour intervals. Again, we observed a difference in N2 and CX12311 animals at both 10% and 21% environmental O_2_ levels. N2 animals were observed to contain fertilized eggs approximately one hour earlier that CX12311 animals (**Figure 3i**). The difference in timing of spermatogenesis and fertilization (2 hours vs 1 hour), potentially reflects the fact that N2 animals produce more sperm before switching to oogenesis.

These experiments suggest that the differences in transcription between N2 and CX12311 could be caused by differences in sexual maturity. We are unable, however, to explain the differences in transcription we observed between 10% and 21% O_2_ as mature sperm was observed at similar times in these different environmental conditions (**Figure 3e**). Potentially the rate of spermatogenesis or expression levels of genes are modified by O_2_ levels that are not reflected in the timing of the presence of mature sperm.

### Derived alleles of *npr-1* and/or *glb-5* increase feeding rates in an O_2_-independent manner

Life-history tradeoffs have been proposed in evolutionary theory to account for the linkage between two different traits. Assuming an individual can acquire a finite amount of energy, the investment of energy into one trait leads to consequential changes in other traits as energy resources are shunted into different directions. For example, artificial selection experiments on early fecundity in *C. elegans* resulted in decreased reproduction late in life (Anderson et al. 2011). The N2 strain seems to violate this tradeoff, as it sexually matures earlier than CX12311, but also produces more eggs later on in life. We measured the size of N2 and CX12311 animals and found that N2 animals were also larger than CX12311 animals at synchronized time points (**Figure 4a**). These observations suggest that the assumption of a fixed energy acquisition for CX12311 and N2 might be violated. This would be consistent with Andersen’s et al’s observation that metabolism genes were upregulated by the derived *npr-1* allele, which they proposed represented differences in food intake (Andersen et al. 2014). It would also be consistent with the role of homologs of *npr-1* in other species. *npr-1* encodes an ortholog to neuropeptide Y receptors, which are reported to regulate feeding behavior in fishes, birds, and mammals (Ando et al. 2001; Lecklin et al. 2002; Matsuda 2009).

**Figure 4.**
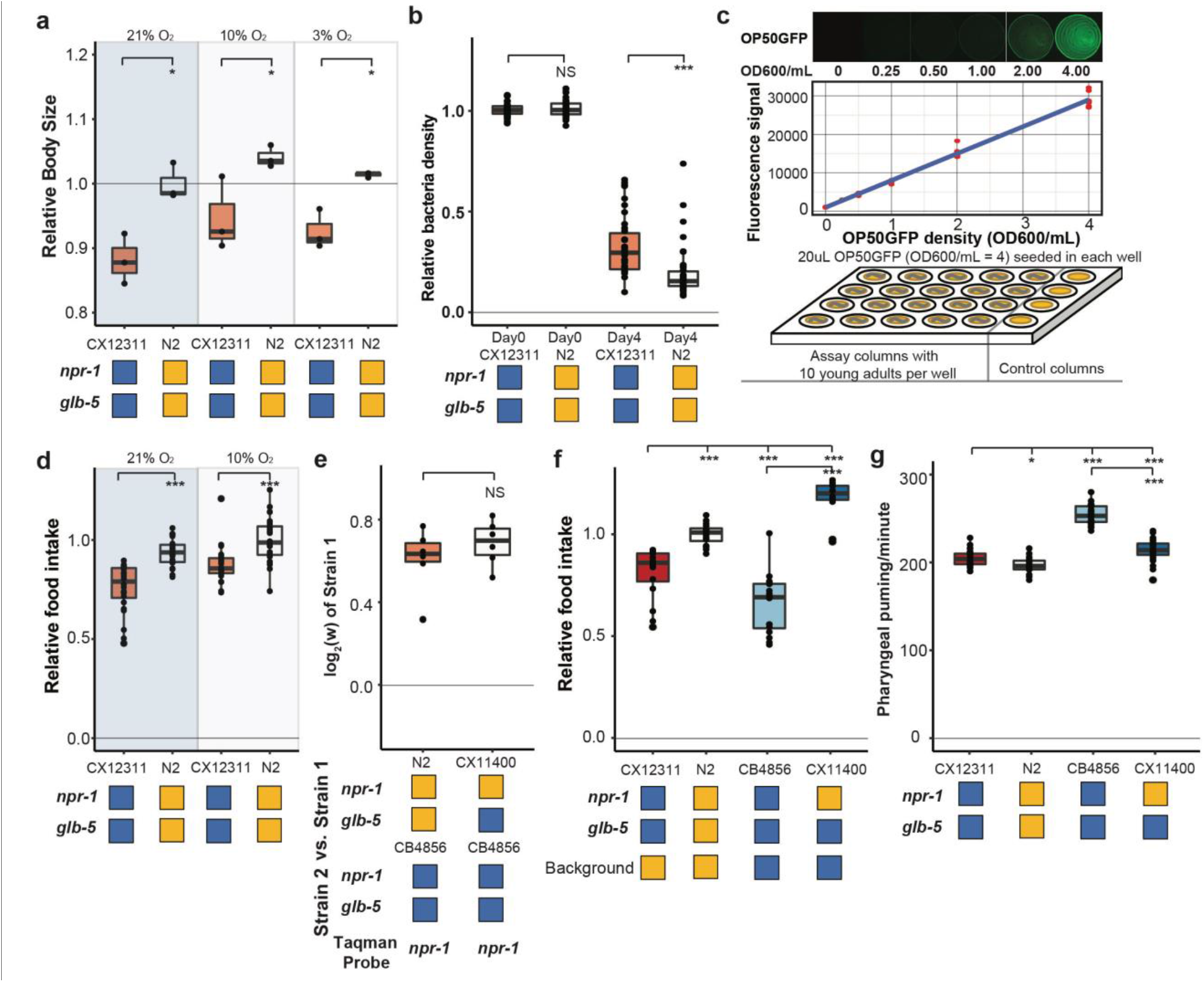
Feeding differences of strains containing derived alleles. **a**. N2 and CX12311 animals were synchronized by hatch-off and allowed to grow at the indicated O_2_ levels for 72 hours. Video recordings were used to estimate the size of the animals. **b**. A previously published liquid, bacterial clearing assay was used to estimate food consumption for the CX12311 and N2 animals. On day 4, N2 animals had consumed more bacteria than CX12311 animals. **c**. To test food consumption on agar plates, we developed a new assay by seeding 24-well agar plates with defined amounts of OP50-GFP bacteria. The number of bacteria on the plate could be estimated using a microplate reader. **d**. N2 animals consumed more food than CX12311 regardless of foraging behaviors. **e**. We tested the effect of the N2 allele of npr-1 in the CB4856 wild strain using the CX11400 NIL strain. Background indicates whether the genomic background is N2 (orange) or CB4856 (blue) **f**. Competition experiments between CB4856 and N2 strains or CB4856 and the CX11400 NIL. **g**. Pharyngeal pumping rates of N2, CB4856 and two NIL strains.

To test this hypothesis, we first utilized a previously described feeding assay to measure the ability of a strain to clear *E. coli* OP50 bacteria from liquid S-media (Gomez-Amaro et al. 2015). In this assay, individual wells are seeded with a defined number of bacteria and 20 worms. Each day, the optical density of each well is measured to estimate the amount of food consumed by the worms. In these conditions, N2 cleared the bacteria faster than CX12311 animals (**Figure 4b**). While these assays supported our hypothesis, liquid media is fundamentally different from the conditions experienced on agar plates, making it difficult to generalize the results from one condition to the other. To this end, we developed a new feeding assay on agar media in 24-well plates. In this assay, each well was seeded with a defined amount of OP50-GFP, which we found could be quantified in a linear manner using a multimode plater reader (**Figure 4c**). When we tested N2 and CX12311 animals in 10% or 21% environmental O_2_ levels, we found N2 consumed more food than CX12311 in both environmental conditions (**Figure 4d**). Interestingly, we also found animals grown in 10% O_2_ also consume more food than animals grown in 21% O_2_. These experiments indicate that N2 animals consume more food than CX12311.

We next decided to test whether the derived allele of *npr-1* could increase the fitness and feeding rate in a different genetic background. We used the CB4856 strain, which has relatively low relative fitness in laboratory conditions (**Figure 2d**), taking advantage of a previously constructed NIL of *npr-1* introgressed from N2 into the CB4856 background (CX11400) (Bendesky et al. 2012). We found that the N2 region surrounding *npr-1* also conferred a fitness advantage in the CB4856 background (**Figure 4e**). The estimated selective coefficients of the derived allele of *npr-1* was higher in the CB4856 background than the N2 background (s = 0.61 vs s = 0.30), potentially due to the lower relative fitness of the CB4856 strain. The food consumption of these strains was consistent with the fitness differences (**Figure 4f**). The derived allele of *npr-1* increased food consumption in both genetic backgrounds but its effect was higher in CB4856.

Food is consumed from the environment by the periodic contraction and relaxation of the pharyngeal muscle which serves to bring material from the environment into the pharynx and filter out bacterial cells (Fang-Yen, Avery, and Samuel 2009). To test whether the increase in food consumption could be explained by an increase in the rate of pumping, we measured the pharyngeal pumping rate of the CX12311, N2, CB4856, and CX11400 strains. The effects of the derived allele of *npr-1* was epistatic with respect to the N2 or CB4856 background. The derived allele decreased the pumping rate in the CB4856 background but had no effect in the N2 background (**Figure 4g**). The effect of the derived allele of *npr-1* is opposite to our expectation and indicates that increased pharyngeal pumping is not responsible for the increased feeding rates. We also measured a number of size parameters of the pharynx but found no obvious differences that could account for the increased food consumption (**Figure S1**). Potentially the pharynx is more efficient at bringing food in from the external environment due to stronger pump strength or more efficient filtering processes.

### Fitness gains of the derived alleles require the URX, AQR, and/or PQR neurons and ascaroside pheromones

We next decided to gain insight into the cellular mechanisms by which *npr-1* and *glb-5* increased fitness of the strains. Previous studies have shown that *npr-1* and *glb-5* regulate social behavior through the URX-RMG neuronal circuit. *glb-5* senses oxygen level changes in the URX oxygen-sensing neuron pair leading to an influx of Ca++ into the cell body (McGrath et al. 2009; Teotonio et al. 2017). The derived allele of *npr-1* inhibits the activity of the RMG hub interneuron which suppresses aerotaxis and social behavior (Laurent et al. 2015; Macosko et al. 2009). The RMG neurons connect to URX and a number of other sensory neurons through gap junctions, which are necessary for foraging behaviors (Jang et al. 2017). URX neurons also integrate O_2_ with internal nutrient reserves (Witham et al. 2016). To test the role of URX in the fitness gains of the *npr-1* and *glb-5* derived alleles, we used the *qaIs2241* integrated cassette that genetically ablates the O_2_ sensing neurons URX, AQR and PQR (Chang et al. 2006). We crossed this cassette into the QG1, CX10774 and CX12311 NILs and repeated the pairwise competition experiments performed in **Figure 1g** using strains that now also contained the *qaIs2241* cassette. In all cases, the relative fitness gains of the derived alleles were decreased by the presence of the neuronal ablation (**Figure 5a**). In one pairwise competition (CX12311 vs QG1), the presence of the *qaIs2241* cassette resulted in the derived allele of *glb-5* decreasing fitness of the animals, suggesting that other *glb-5* expressing sensory neurons (BAG, ADF, or ASG) can modulate fitness of the animals. These experiments suggest that the derived alleles either activate or disinhibit the URX, AQR, and or PQR neurons which leads to increases in fitness.

**Figure 5.**
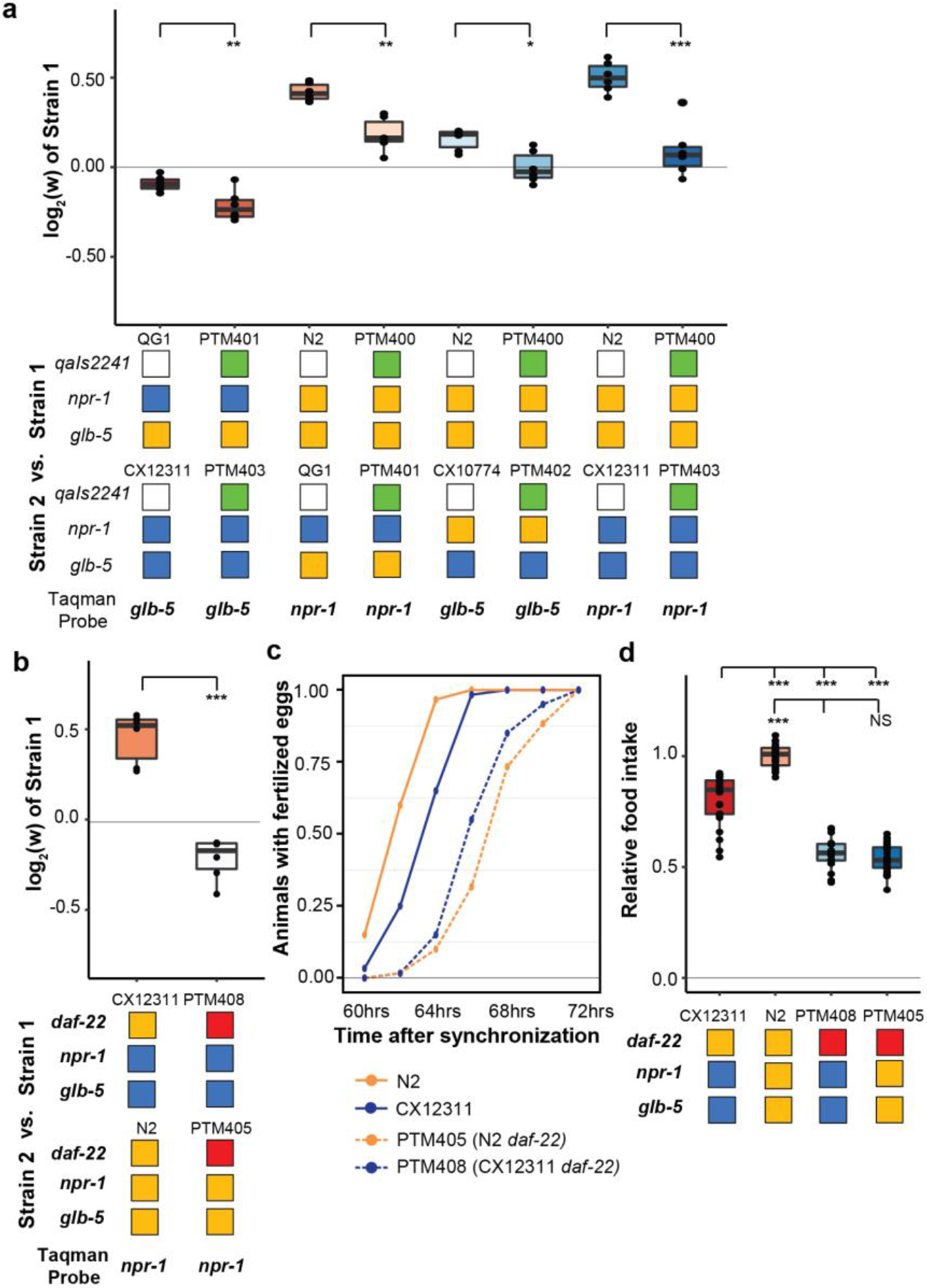
O_2_-sensing neurons and a pheromone biosynthesis enzyme contribute to fitness differences of N2 and CX12311. **a**. Competition experiments between indicated strains. *qaIs2241* is an integrated genetic cassette that ablates the URX, AQR, and PQR neurons. Green indicates the presence of the cassette. **b**. Competition experiments between indicated strains. *daf-22* encodes a sterol carrier protein, which is required for biosynthesis of most ascaroside pheromones. Red indicates the strain contain a deletion that spans the gene. **c**. Number of animals that carry fertilized eggs at the indicated timepoints. **d**. On plate feeding assays of the indicated strains.

We also decided to test whether ascaroside pheromones were necessary for the fitness differences between N2 and CX12311. Nematodes release a number of ascaroside molecules, which are in turn sensed by a distributed neural circuit that integrates and modifies a number of behavioral and developmental phenotypes (Butcher 2017; Ludewig and Schroeder 2013). There are a few reasons to think that ascaroside pheromones might be involved in the fitness gains of the N2 strain. First, work by Andersen et. al indicated that population density directly impacts lifetime fecundity and adult body length differences between N2 and CB4856 strains (Andersen et al. 2014). Second, our previous studies of *C. elegans* domestication to liquid cultures has found that pheromone signaling was modified by fixed genetic changes (Large et al. 2016; McGrath et al. 2011). Finally, the derived alleles of *npr-1* and *glb-5* have been shown to modify pheromone valence in a variety of contexts (Fenk and de Bono 2017; Jang et al. 2012; Macosko et al. 2009; Oda, Toyoshima, and de Bono 2017). To test the role of ascaroside pheromones, we followed previous publications using a genetic knockout of the *daf-22* gene, which encodes a peroxisomal enzyme required for the biosynthesis of *C. elegans* pheromones (Butcher et al. 2009) and accumulation of lipid droplets (Zhang et al. 2010), using CRISPR-Cas9 enabled genome editing to create a large deletion of *daf-22* in the N2 strain, which was then crossed to the CX12311 background. Competition experiments demonstrated that *daf-22* was necessary for the fitness advantage of derived *npr-1* and *glb-5* alleles (**Figure 5b**). In addition, *daf-22* was necessary for the faster sexual maturity (**Figure 5c**) and increased food intake (**Figure 5d**) of the N2 strain compared to CX12311. These data suggest that *npr-1* and *glb-5* reprogram pheromone responses resulting in increased sexual maturity and ability to consume food.

## Discussion

In this report, we studied the fitness consequences of two derived alleles that arose and fixed in the N2 strain after isolation from the wild. We find that both alleles can be adaptive, with selective coefficients that are larger than many characterized beneficial alleles from other species. These results are consistent with the derived alleles spreading through the ancestral N2 populations due to positive selection. If this was true, it would suggest that the derived allele of *npr-1* arose first, as the derived *glb-5* allele is only beneficial in this derived genetic background. However, the demographic history and laboratory environment of how N2 was grown at the time these alleles arose is largely lost (Sterken et al. 2015). The exact laboratory growth conditions (liquid axenic vs. solid media), transfer processes (picking vs. chunking) and effective population sizes (between 4 and 1000) used to propagate a *C. elegans* strain is incredibly variable. It is likely that the evolutionary forces responsible for the fixation of these alleles will remain lost to history.

Nevertheless, the ability of positive selection to act upon the derived *npr-1* allele can be observed in current experiments. A recent example is provided by Noble and colleagues, who created a large mapping population between sixteen parental strains (including N2 and CB4856) to create a large panel recombinant inbred lines (RILs) (Noble et al. 2017). During the outcrossing phase of construction, the N2 allele of *npr-1* spread through the population to fixation, consistent with its dominant action and the strong selective advantage of this allele. Potentially, variation in *npr-1* affected allele frequencies of unlinked loci as well. For example, an excess of CB4856 haplotypes was observed in the RILs, suggesting that CB4856 haplotypes were more likely to contain beneficial alleles. Our measurements of the relative fitness of the CB4856 strain, however, creates an apparent paradox, as CB4856 was one of the least fit strains among the wild strains we tested (**Figure 2d**). Potentially, epistatic interactions between CB4856 alleles and the derived allele of *npr-1* could help resolve this; the effect of *npr-1* on food intake and fitness is higher in the CB4856 background (**Figure 4d,e**). Differences in effect size of a focal allele in different genetic backgrounds is considered evidence for the existence of epistasis (Gibson and Dworkin 2004). Potentially, the presence of laboratory-derived alleles in mapping populations will skew not only the allele frequencies of these beneficial alleles, but also natural genetic variants that interact epistatically with them.

Evolution of behavioral traits is one strategy for animals to respond to a new environment. The identification of a polymorphism in *npr-1* has served as an example of how behavioral variation can arise from genetic variation. However, our work suggests that the behavioral changes of N2 are not sufficient for explaining its fitness gains. Rather, we propose that changes to food intake, sexual maturity, and fecundity are more important. One unresolved question is why wild strains do not eat as much food as the N2 strain? We believe there must be some sort of tradeoff – either energetically or developmentally – that makes the derived mutation unfavorable in their natural environments. Mechanistic understanding of the energetic forces necessary for *C. elegans* to bring food into their pharynx is lacking. In fact, pharyngeal pumping rates are often used as proxies for food intake, which we have shown here are unrelated to the amount of food consumed. Potentially the thick slurry of food in laboratory plates is completely different biophysically from the mixed bacterial species encountered on rotting material in the wild. Alternatively, differences in feeding behaviors unrelated to social/solitary behaviors might also mediate the differences in food intake.

The changes to fitness and feeding rate appear to be mediated by the nervous system, potentially as a result to changes in pheromone responses. Primer pheromones regulate physiology and have been shown to influence feeding and metabolism in *C. elegans* (Hussey et al. 2017) and other species (Botzer et al. 1991). Modification of pheromone responses might represent a common strategy in nematode evolution, both in the laboratory and in natural populations. Large scale identification of beneficial alleles that confer fitness advantages has largely occurred in unicellular organisms (Herron and Doebeli 2013; Kvitek and Sherlock 2013; Venkataram et al. 2016); similar high-throughput experiments in nematodes should be illuminating in this regard.

Our work underscores issues with growing organisms in the laboratory for multiple generations. Despite the attempts of researchers to create fertile conditions for nematodes to grow in, we found a large difference in relative fitness between different strains of *C. elegans* when competed in the laboratory. Natural genetic variation and *de novo* variation both result in fitness differences that selection can act on. Experimenters using wild strains of nematodes must take care in designing experiments to account for this, especially in wild strains with lower initial fitness levels. We believe that the laboratory selection pressures we characterized here will generalize to other invertebrate and vertebrate animals. If so, the behaviors and physiology of these animals will also be modified over generations of growth. Our work suggests that not only will the traits that confer fitness advantages be modified, but potentially additional traits due to the pleiotropic actions of many genes, and relaxed stabilizing selection on traits in laboratory conditions.

## Acknowledgements

Some strains were provided by the Andersen lab, the Bargmann lab, and the CGC, which is funded by NIH Office of Research Infrastructure Programs (P40 OD010440). We thank Greg Gibson’s lab for assistance in Droplet Digital PCR. We are also grateful to Erik Andersen, Cori Bargmann, Levi Morran, Annalise Paaby, and members of the McGrath lab for comments on the manuscript.

## Methods

### Strains

The following strains were used in this study:

Wild strains: N2; CB4856; DL238; JU258; JU775; MY16; MY23; CX11314; LKC34; ED3017; JT11398; EG4725. The N2 strain originated from the Bargmann lab (The Rockefeller University). The remaining eleven wild strains came from the *Caenorhabditis elegans* Natural Diversity Resource (Cook et al. 2017).

Barcoded strains: PTM229 *dpy-10 (kah82)II;* PTM288 *dpy-10 (kah83)II kyIR1(V, CB4856>N2) qgIR1(X, CB4856>N2)*; PTM289 *dpy-10 (kah84)II kyIR1(V, CB4856>N2) qgIR1(X, CB4856>N2)*; The barcoded strains were generated using previously published reagents for modifying the *dpy-10* gene (Arribere et al. 2014). Two modified repair oligos with the following sequence were used to edit silent mutations into the 90^th^ (Thr) or 92^nd^ amino acid (Arg):

*dpy-10* 90^th^ silent mutation:

5’-

CACTTGAACTTCAATACGGCAAGATGAGAATGACTGGAAACCGTACTGCTCGTGGTGCCTATGGTAGCGGAGCTTCACATGGCTTCAGACCAACAGCCTAT-3’

*dpy-10* 92^nd^ silent mutation:

5’-

CACTTGAACTTCAATACGGCAAGATGAGAATGACTGGAAACCGTACCGCTCGCGGTGCCTATGGTAGCGGAGCTTCACATGGCTTCAGACCAACAGCCTAT-3’

The microinjection mix was: 50ng/uL *Peft3::Cas9*, 25ng/uL *dpy-10* sgRNA, 500nM *dpy-10(cn64)* repair oligo, and one of the 500nM *dpy-10(90/92)* repair oligo. This mix was injected into N2 or CX12311 and so-called “jackpot broods” were identified by the presence of a large number of F1 animals with the roller phenotype. From these plates, wildtype animals were singled and genotyped using Sanger-sequencing. *kah82* and *kah83* contain the 90^th^ Thr silent mutation (ACC -> ACT). *kah84* contains the 92^nd^ Arg silent mutation (CGT -> CGC).

Near isogenic lines: CX12311 *kyIR1(V, CB4856>N2) qgIR1(X, CB4856>N2);* QG1 *qgIR1(X, CB4856>N2);* CX10774 *kyIR1(V, CB4856>N2);* CX11400 *kyIR9(X, N2>CB4856)*. These strains were originally described in previous studies (Bendesky et al. 2012; Bernstein and Rockman 2016; McGrath et al. 2009; McGrath et al. 2011).

*npr-1* loss of function: CX4148 *npr-1(ky13)X;* DA609 *npr-1(ad609)X;* These strains were previously described (de Bono and Bargmann 1998)

URX, AQR, PQR genetic ablation strains: qals2241 *[Pgcy-35::GFP Pgcy-36::egl-1 lin15+]* is an integrated transgene that genetically ablates URX, AQR, and PQR neurons (Chang et al. 2006). This transgene was crossed into a number of introgressed regions using standard genetic techniques. CX7102 *qaIs2241X;* PTM400 *qals2241X;* PTM401 *qgIR1(X, CB4856>N2) qaIs2241X;* PTM402 *kyIR1 (V, CB4856>N2) qaIs2241X*; PTM403 *kyIR1(V, CB4856>N2) qgIR1(X, CB4856>N2) qaIs2241X*;

*daf-22* strains: *daf-22(kah8)II* is a *daf-22* gene disruption made by CRISPR/Cas9 genome editing (Large et al. 2016). This transgene was crossed into a number of introgressed regions using standard genetic techniques. PTM95 *daf-22(kah8)II kyIR1(V, CB4856>N2) qgIR1(X, CB4856>N2);* PTM404 *daf-22(kah8)II dpy10(kah83)II;* PTM405 *daf-22(kah8)II;* PTM408 *daf-22(kah8)II kyIR1(V, CB4856>N2) qgIR1(X, CB4856>N2)*.

### Growth conditions

Animals were grown following standard conditions. With exceptions listed below, animals were cultivated on modified nematode growth medium (NGM) plates containing 2% agar seeded with 200 ul of an overnight culture of the *E. coli* strain OP50 in an incubator set at 20°C. Strains were grown for at least three generations without starvation before any assays were conducted. For assays manipulating the environmental O_2_ levels, animals were grown inside a BioSpherix C474 chamber using a BioSpherix C21 single chamber controller to control ambient O_2_ levels. For these assays, animals were not grown in temperature incubators, and the room temperature was typically kept ~21 °C. For competition experiments on non-burrowing plates, 1.25% agarose and 0.75% agar replaced the agar concentrations of normal growth plates. To create uniform lawns, OP50 bacteria was poured onto plates to cover the entire surface area of the plate and then poured off.

### Pairwise fitness measurements

Competition experiments were performed essentially as in the previous study (Large et al. 2016). Ten L4 stage animals from each strain were picked onto 9cm NGM plates seeded with 300uL of an overnight *E. coli* OP50 culture and incubated at room temperature for three days. After five days, animals were transferred to an identically-prepared NGM plate and then subsequently transferred every four days for five to seven generations. For transfers, animals were washed off from the test plates using M9 buffer and collected into 1.5mL centrifuge tube. The animals were mixed by inversion and allowed to stand for approximately one minute to settle adult animals. 50uL of the supernatant containing ~1000-2000 L1-L2 animals were seeded on next plates. The remaining animals were concentrated and placed in a −80°C freezer for future genomic DNA isolation. Genomic DNA was collected from every odd generation using a Zymo DNA isolation kit (D4071).

To quantify the relative proportion of each strain, we used a digital PCR based approach using a custom TaqMan probe (Applied Biosciences). Genomic DNA was digested with EcoRI for 30 min at 37 ^°^C. The digested products were purified using a Zymo DNA cleanup kit (D4064) and diluted to ~1ng/uL for the following Taqman assay. Four TaqMan probes were designed using ABI custom software that targeted the *dpy-10 (kah82), dpy-10 (kah84), npr-1(g320)*, or SNP WBVar00209467 in *glb-5*. These probes were validated using defined concentrations of DNA from animals containing each allele. The Taqman digital PCR assays were performed using a Biorad QX200 digital PCR machine with standard probe absolute quantification protocol. The relative allele proportion was calculated for each DNA sample using count number of the droplet with fluorescence signal (equation 1). To calculate the relative fitness of the two strains using three to four measurements of relative fitness, we used linear regression to fit this data to a one-locus generic selection model (equation 2 and 3), assuming one generation per transfer.

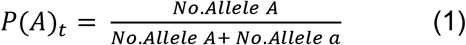

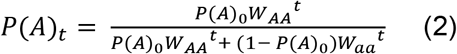

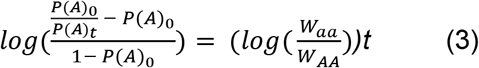

### Aerotaxis assays

To measure bordering rates, two -week old NGM plates were removed from a 4°C cold room, seeded with 200uL of *E. coli* OP50 and incubated for two days at room temperature. 150 adult animals were picked onto these assay plates and placed in either a 20 °C incubator or a BioSpherix chamber for 3 hours. Bordering behavior was quantified using a dissecting microscope by identifying animals whose whole body resided within 1mm of the border of the bacteria lawn.

### Transcriptome analysis

N2 and CX12311 L4 hermaphrodites were picked to fresh agar plates. Their adult progeny was synchronized using alkaline-bleach to isolate eggs. These eggs were washed three times using M9 buffer and placed on a tube roller overnight to allow eggs to hatch. About 400 L1 animals were placed on NGM agar plates seeded with *E. coli* OP50 and incubated in a BioSpherix chamber set at 10% O_2_ or 21% O_2_ levels for 48 hours. The ~L4 stage animals were washed off and used for standard Trizol RNA isolation. Replicates were performed on different days. The RNA libraries for next-generation sequencing were prepared using an Illumina TruSeq Stranded mRNA kit (20020595) following its standard protocol. These libraries were sequenced using an Illumina NextSeq 500 platform. Reads were aligned using HISAT2 using default parameters for pair-end sequencing. Transcripts abundance was calculated using HTseq and then used as inputs for the SARTools (Varet et al. 2016). Within this R package, edgeR is used for normalization and differential analysis. N2 cultured at 21% O2 is treated as wild type (Chen, Lun, and Smyth 2014). The genes show different expression (log2(fold) > 1 or log2(fold) < −1, FDR adjusted p-value < 0.01) were selected to perform Hierarchical Cluster analysis, and Principal Component analysis. Sequencing reads were uploaded to the SRA under PRJNA437304.

### Feeding rate, pharyngeal pumping rate, and pharyngeal size assays

The 24 well-plates were prepared by pipetting 0.75mL NGM agar contain 25uM FUDR and 1x Antibiotic-Antimycotic (ThermoFisher 15240062) to each well. The fresh prepared plates were placed in fume hood and dried with air flow for 1.5 hours. 20uL of freshly cultured OD600 of 4.0 (CFU ~ 3.2×10^9^/mL) *E. coli* OP50-GFP(pFPV25.1) were seeded in the center of each well. Animals were synchronized using alkaline-bleach. The eggs were washed by M9 buffer for three times and rotating on tube roller overnight to allow eggs to hatch. About 200 L1 animals were placed on NGM agar plates seeded with *E. coli* OP50 and cultivate at 20 °C or BioSpherix chamber at 21 °C for 50 hours. Ten animals (Late L4 stage or young adult) were transferred to each well of the first 5 columns of the feeding rates assay 24 well-plates. The rest 4 wells were used to measure the GFP signal degradation during the feeding rates assay. After place animals on the feeding rates assay plates, the fluorescence signal of OP50-GFP from each well was quantified by area scanning protocol using Bio Tek Synergy H4 multimode plate reader at 6mm height as the starting time point. The 24-well plates were then incubated in incubator or BioSpherix chamber for 18 hours and the fluorescence signal were quantified again as the ending time point. The bacteria amount at end time point from each well was normalized using the fluorescence signal degradation amount of control wells. The normalization was performed using the equation as below:

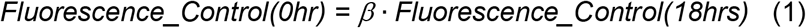

All of the signals from control wells were used to do linear regression and estimate coefficient β. The actual amounts of bacteria at 18hrs for each test is:

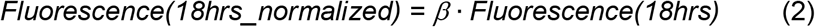

The food consumption for each well was calculated by:

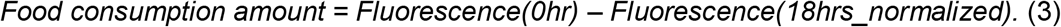

### Pharyngeal pumping and size assays

Animals were synchronized using alkaline-bleach. The eggs were washed by M9 buffer for three times and rotating on tube roller overnight to allow eggs to hatch. About 200 L1 animals were placed on NGM agar plates seeded with *E. coli* OP50 and cultivated at 20°C for 72 hours. In the pharyngeal pumping rates assays, the pharynges of ten young adult animals (72 hours after place L1 on NGM agar plate) were observed for 30 seconds each in three separate trails. To measure the pharyngeal size, young adult animals were placed onto agar pad and immobilized by 25mM NaN_3_. To each strain, pharyngeal sizes of 30 animals from three different plates were imaged under 40x objective lens using z-stack DIC microscope. The diameter of pharyngeal metacorpus, diameter of terminal bulb diameter, procorpus length, and isthmus length were measured by ImageJ software.

### Reproductive timing and growth assays

To measure reproductive timing, animals were synchronized by picking ten adult animals onto an NGM plate, allowing them to lay eggs for two hours, and then removing the adult animals from the plate. These offspring were then monitored using a 12x dissecting microscope at indicated time points to count the number of animals with oocytes and fertilized eggs in their uterus. A subset of these animals was washed off at indicated time points and fixed in 95% ethanol and stained with DAPI. Each spermathecal was imaged by z-stack fluorescence microscopy using a 100x lens to determine whether spermatogenesis had started or two count the number of sperm produced by the hermaphrodite.

Reproductive rate and body size measurements were measured as described previously (Large et al. 2016).

### Statistics

To assess statistical significance, we performed one-way ANOVA tests followed by Tukey’s honest significant difference test to correct for multiple comparisons for data presented in 1f, 1g, 1h, 2b, 2c, 2d, 2e, 4f, 4g, and 5d. To test pairwise comparisons presented in Figures 2a, 3f, 3h, 4a, 4b, 4d, 4e, 5a, and 5b, we also used the Wilcoxon-Mann-Whitney nonparametric test. The Friedman test was used to compare the reproductive timing assays shown in 3e, 3i, 5c.

## References

Alfred, J., and I. T. Baldwin. 2015. ’New opportunities at the wild frontier’, Elife, 4.

Andersen, E. C., J. S. Bloom, J. P. Gerke, and L. Kruglyak. 2014. ’A variant in the neuropeptide receptor npr-1 is a major determinant of Caenorhabditis elegans growth and physiology’, PLoS Genet, 10: e1004156.

Anderson, J. L., R. M. Reynolds, L. T. Morran, J. Tolman-Thompson, and P. C. Phillips. 2011. ’Experimental evolution reveals antagonistic pleiotropy in reproductive timing but not life span in Caenorhabditis elegans’, J Gerontol A Biol Sci Med Sci, 66: 1300–8.

Ando, R., S. I. Kawakami, T. Bungo, A. Ohgushi, T. Takagi, D. M. Denbow, and M. Furuse. 2001. ’Feeding responses to several neuropeptide Y receptor agonists in the neonatal chick’, Eur J Pharmacol, 427: 53–9.

Arribere, J. A., R. T. Bell, B. X. Fu, K. L. Artiles, P. S. Hartman, and A. Z. Fire. 2014. ’Efficient marker-free recovery of custom genetic modifications with CRISPR/Cas9 in Caenorhabditis elegans’, Genetics, 198: 837–46.

Bendesky, A., J. Pitts, M. V. Rockman, W. C. Chen, M. W. Tan, L. Kruglyak, and C. I. Bargmann. 2012. ’Long-range regulatory polymorphisms affecting a GABA receptor constitute a quantitative trait locus (QTL) for social behavior in Caenorhabditis elegans’, PLoS Genet, 8: e1003157.

Bernstein, M. R., and M. V. Rockman. 2016. ’Fine-Scale Crossover Rate Variation on the Caenorhabditis elegans X Chromosome’, G3 (Bethesda), 6: 1767–76.

Bersaglieri, T., P. C. Sabeti, N. Patterson, T. Vanderploeg, S. F. Schaffner, J. A. Drake, M. Rhodes, D. E. Reich, and J. N. Hirschhorn. 2004. ’Genetic signatures of strong recent positive selection at the lactase gene’, Am J Hum Genet, 74: 1111–20.

Boeck, M. E., C. Huynh, L. Gevirtzman, O. A. Thompson, G. Wang, D. M. Kasper, V. Reinke, L. W. Hillier, and R. H. Waterston. 2016. ’The time-resolved transcriptome of C. elegans’, Genome Res, 26: 1441–50.

Botzer, D., S. Blumberg, I. Ziv, and A. J. Susswein. 1991. ’Common regulation of feeding and mating in Aplysia fasciata: pheromones released by mating and by egg cordons increase feeding behavior’, Behav Neural Biol, 56: 251–61.

Brenner, S. 1974. ’The genetics of Caenorhabditis elegans’, Genetics, 77: 71–94.

Butcher, R. A. 2017. ’Decoding chemical communication in nematodes’, Nat Prod Rep, 34: 472–77.

Butcher, R. A., J. R. Ragains, W. Li, G. Ruvkun, J. Clardy, and H. Y. Mak. 2009. ’Biosynthesis of the Caenorhabditis elegans dauer pheromone’, Proc Natl Acad Sci U S A, 106: 1875–9.

Chang, A. J., N. Chronis, D. S. Karow, M. A. Marletta, and C. I. Bargmann. 2006. ’A distributed chemosensory circuit for oxygen preference in C. elegans’, PLoS Biol, 4: e274.

Chen, Yunshun, Aaron TL Lun, and Gordon K Smyth. 2014. ’Differential expression analysis of complex RNA-seq experiments using edgeR.’ in, Statistical analysis of next generation sequencing data (Springer).

Cheung, B. H., M. Cohen, C. Rogers, O. Albayram, and M. de Bono. 2005. ’Experience-dependent modulation of C. elegans behavior by ambient oxygen’, Curr Biol, 15: 905–17.

Coates, J. C., and M. de Bono. 2002. ’Antagonistic pathways in neurons exposed to body fluid regulate social feeding in Caenorhabditis elegans’, Nature, 419: 925–9.

Cook, D. E., S. Zdraljevic, J. P. Roberts, and E. C. Andersen. 2017. ’CeNDR, the Caenorhabditis elegans natural diversity resource’, Nucleic Acids Res, 45: D650–D57.

de Bono, M., and C. I. Bargmann. 1998. ’Natural variation in a neuropeptide Y receptor homolog modifies social behavior and food response in C. elegans’, Cell, 94: 679–89.

Duveau, F., and M. A. Felix. 2012. ’Role of pleiotropy in the evolution of a cryptic developmental variation in Caenorhabditis elegans’, PLoS Biol, 10: e1001230.

Evans, K. S., Y. Zhao, S. C. Brady, L. Long, P. T. McGrath, and E. C. Andersen. 2017. ’Correlations of Genotype with Climate Parameters Suggest Caenorhabditis elegans Niche Adaptations’, G3 (Bethesda), 7: 289–98.

Fang-Yen, C., L. Avery, and A. D. Samuel. 2009. ’Two size-selective mechanisms specifically trap bacteria-sized food particles in Caenorhabditis elegans’, Proc Natl Acad Sci U S A, 106: 20093–6.

Fenk, L. A., and M. de Bono. 2017. ’Memory of recent oxygen experience switches pheromone valence in Caenorhabditis elegans’, Proc Natl Acad Sci U S A, 114: 4195–200.

Fisher, K. J., and G. I. Lang. 2016. ’Experimental evolution in fungi: An untapped resource’, Fungal Genet Biol, 94: 88–94.

Gibson, G., and I. Dworkin. 2004. ’Uncovering cryptic genetic variation’, Nat Rev Genet, 5: 681–90.

Gladfelter, A. S. 2015. ’How nontraditional model systems can save us’, Mol Biol Cell, 26: 3687–9.

Gloria-Soria, A., and R. B. Azevedo. 2008. ’npr-1 Regulates foraging and dispersal strategies in Caenorhabditis elegans’, Curr Biol, 18: 1694–9.

Goldstein, B., and N. King. 2016. ’The Future of Cell Biology: Emerging Model Organisms’, Trends Cell Biol, 26: 818–24.

Gomez-Amaro, R. L., E. R. Valentine, M. Carretero, S. E. LeBoeuf, S. Rangaraju, C. D. Broaddus, G. M. Solis, J. R. Williamson, and M. Petrascheck. 2015. ’Measuring Food Intake and Nutrient Absorption in Caenorhabditis elegans’, Genetics, 200: 443–54.

Goto, T., A. Tanave, K. Moriwaki, T. Shiroishi, and T. Koide. 2013. ’Selection for reluctance to avoid humans during the domestication of mice’, Genes Brain Behav, 12: 760–70.

Gould, S. J., and R. C. Lewontin. 1979. ’The spandrels of San Marco and the Panglossian paradigm: a critique of the adaptationist programme’, Proc R Soc Lond B Biol Sci, 205: 581–98.

Gray, J. M., D. S. Karow, H. Lu, A. J. Chang, J. S. Chang, R. E. Ellis, M. A. Marletta, and C. I. Bargmann. 2004. ’Oxygen sensation and social feeding mediated by a C. elegans guanylate cyclase homologue’, Nature, 430: 317–22.

Greene, J. S., M. Brown, M. Dobosiewicz, I. G. Ishida, E. Z. Macosko, X. Zhang, R. A. Butcher, D. J. Cline, P. T. McGrath, and C. I. Bargmann. 2016. ’Balancing selection shapes density-dependent foraging behaviour’, Nature, 539: 254–58.

Herron, M. D., and M. Doebeli. 2013. ’Parallel evolutionary dynamics of adaptive diversification in Escherichia coli’, PLoS Biol, 11: e1001490.

Hubbard, E. J., and D. Greenstein. 2005. ’Introduction to the germ line’, WormBook: 1–4.

Hussey, R., J. Stieglitz, J. Mesgarzadeh, T. T. Locke, Y. K. Zhang, F. C. Schroeder, and S. Srinivasan. 2017. ’Pheromone-sensing neurons regulate peripheral lipid metabolism in Caenorhabditis elegans’, PLoS Genet, 13: e1006806.

Jang, H., K. Kim, S. J. Neal, E. Macosko, D. Kim, R. A. Butcher, D. M. Zeiger, C. I. Bargmann, and P. Sengupta. 2012. ’Neuromodulatory state and sex specify alternative behaviors through antagonistic synaptic pathways in C. elegans’, Neuron, 75: 585–92.

Jang, H., S. Levy, S. W. Flavell, F. Mende, R. Latham, M. Zimmer, and C. I. Bargmann. 2017. ’Dissection of neuronal gap junction circuits that regulate social behavior in Caenorhabditis elegans’, Proc Natl Acad Sci U S A, 114: E1263–E72.

Kasahara, T., K. Abe, K. Mekada, A. Yoshiki, and T. Kato. 2010. ’Genetic variation of melatonin productivity in laboratory mice under domestication’, Proc Natl Acad Sci U S A, 107: 6412–7.

Kvitek, D. J., and G. Sherlock. 2013. ’Whole genome, whole population sequencing reveals that loss of signaling networks is the major adaptive strategy in a constant environment’, PLoS Genet, 9: e1003972.

Large, E. E., R. Padmanabhan, K. L. Watkins, R. F. Campbell, W. Xu, and P. T. McGrath. 2017. ’Modeling of a negative feedback mechanism explains antagonistic pleiotropy in reproduction in domesticated Caenorhabditis elegans strains’, PLoS Genet, 13: e1006769.

Large, E. E., W. Xu, Y. Zhao, S. C. Brady, L. Long, R. A. Butcher, E. C. Andersen, and P. T. McGrath. 2016. ’Selection on a Subunit of the NURF Chromatin Remodeler Modifies Life History Traits in a Domesticated Strain of Caenorhabditis elegans’, PLoS Genet, 12: e1006219.

Laurent, P., Z. Soltesz, G. M. Nelson, C. Chen, F. Arellano-Carbajal, E. Levy, and M. de Bono. 2015. ’Decoding a neural circuit controlling global animal state in C. elegans’, Elife, 4.

Lecklin, A., I. Lundell, L. Paananen, J. E. Wikberg, P. T. Mannisto, and D. Larhammar. 2002. ’Receptor subtypes Y1 and Y5 mediate neuropeptide Y induced feeding in the guinea-pig’, Br J Pharmacol, 135: 2029–37.

Lenski, R. E. 2017. ’Experimental evolution and the dynamics of adaptation and genome evolution in microbial populations’, ISME J, 11: 2181–94.

Li, W. H. 1975. ’The first arrival time and mean age of a deleterious mutant gene in a finite population’, Am J Hum Genet, 27: 274–86.

Ludewig, A. H., and F. C. Schroeder. 2013. ’Ascaroside signaling in C. elegans’, WormBook: 1–22.

Macosko, E. Z., N. Pokala, E. H. Feinberg, S. H. Chalasani, R. A. Butcher, J. Clardy, and C. I. Bargmann. 2009. ’A hub-and-spoke circuit drives pheromone attraction and social behaviour in C. elegans’, Nature, 458: 1171–5.

Marks, M. E., C. M. Castro-Rojas, C. Teiling, L. Du, V. Kapatral, T. L. Walunas, and S. Crosson. 2010. ’The genetic basis of laboratory adaptation in Caulobacter crescentus’, J Bacteriol, 192: 3678–88.

Matsuda, K. 2009. ’Recent advances in the regulation of feeding behavior by neuropeptides in fish’, Ann N Y Acad Sci, 1163: 241–50.

McGrath, P. T., M. V. Rockman, M. Zimmer, H. Jang, E. Z. Macosko, L. Kruglyak, and C. I. Bargmann. 2009. ’Quantitative mapping of a digenic behavioral trait implicates globin variation in C. elegans sensory behaviors’, Neuron, 61: 692–9.

McGrath, P. T., Y. Xu, M. Ailion, J. L. Garrison, R. A. Butcher, and C. I. Bargmann. 2011. ’Parallel evolution of domesticated Caenorhabditis species targets pheromone receptor genes’, Nature, 477: 321–5.

Noble, L. M., I. Chelo, T. Guzella, B. Afonso, D. D. Riccardi, P. Ammerman, A. Dayarian, S. Carvalho, A. Crist, A. Pino-Querido, B. Shraiman, M. V. Rockman, and H. Teotonio. 2017. ’Polygenicity and Epistasis Underlie Fitness-Proximal Traits in the Caenorhabditis elegans Multiparental Experimental Evolution (CeMEE) Panel’, Genetics, 207: 1663–85.

Oda, S., Y. Toyoshima, and M. de Bono. 2017. ’Modulation of sensory information processing by a neuroglobin in Caenorhabditis elegans’, Proc Natl Acad Sci U S A, 114: E4658–E65.

Orozco-terWengel, P., M. Kapun, V. Nolte, R. Kofler, T. Flatt, and C. Schlotterer. 2012. ’Adaptation of Drosophila to a novel laboratory environment reveals temporally heterogeneous trajectories of selected alleles’, Mol Ecol, 21: 4931–41.

Persson, A., E. Gross, P. Laurent, K. E. Busch, H. Bretes, and M. de Bono. 2009. ’Natural variation in a neural globin tunes oxygen sensing in wild Caenorhabditis elegans’, Nature, 458: 1030–3.

Reddy, K. C., E. C. Andersen, L. Kruglyak, and D. H. Kim. 2009. ’A polymorphism in npr-1 is a behavioral determinant of pathogen susceptibility in C. elegans’, Science, 323: 382–4.

Russell, J. J., J. A. Theriot, P. Sood, W. F. Marshall, L. F. Landweber, L. Fritz-Laylin, J. K. Polka, S. Oliferenko, T. Gerbich, A. Gladfelter, J. Umen, M. Bezanilla, M. A. Lancaster, S. He, M. C. Gibson, B. Goldstein, E. M. Tanaka, C. K. Hu, and A. Brunet. 2017. ’Non-model model organisms’, BMC Biol, 15: 55.

Stanley, C. E., Jr., and R. J. Kulathinal. 2016. ’Genomic signatures of domestication on neurogenetic genes in Drosophila melanogaster’, BMC Evol Biol, 16: 6.

Sterken, M. G., L. B. Snoek, J. E. Kammenga, and E. C. Andersen. 2015. ’The laboratory domestication of Caenorhabditis elegans’, Trends Genet, 31: 224–31.

Teotonio, H., S. Estes, P. C. Phillips, and C. F. Baer. 2017. ’Experimental Evolution with Caenorhabditis Nematodes’, Genetics, 206: 691–716.

Varet, Hugo, Loraine Brillet-Guéguen, Jean-Yves Coppée, and Marie-Agnès Dillies. 2016. ’SARTools: a DESeq2-and edgeR-based R pipeline for comprehensive differential analysis of RNA-Seq data’, PLoS One, 11: e0157022.

Venkataram, S., B. Dunn, Y. Li, A. Agarwala, J. Chang, E. R. Ebel, K. Geiler-Samerotte, L. Herissant, J. R. Blundell, S. F. Levy, D. S. Fisher, G. Sherlock, and D. A. Petrov. 2016. ’Development of a Comprehensive Genotype-to-Fitness Map of Adaptation-Driving Mutations in Yeast’, Cell, 166: 1585–96 e22.

Weber, K. P., S. De, I. Kozarewa, D. J. Turner, M. M. Babu, and M. de Bono. 2010. ’Whole genome sequencing highlights genetic changes associated with laboratory domestication of C. elegans’, PLoS One, 5: e13922.

Witham, E., C. Comunian, H. Ratanpal, S. Skora, M. Zimmer, and S. Srinivasan. 2016. ’C. elegans Body Cavity Neurons Are Homeostatic Sensors that Integrate Fluctuations in Oxygen Availability and Internal Nutrient Reserves’, Cell Rep, 14: 1641–54.

Yvert, G., R. B. Brem, J. Whittle, J. M. Akey, E. Foss, E. N. Smith, R. Mackelprang, and L. Kruglyak. 2003. ’Trans-acting regulatory variation in Saccharomyces cerevisiae and the role of transcription factors’, Nat Genet, 35: 57–64.

Zhang, S. O., A. C. Box, N. Xu, J. Le Men, J. Yu, F. Guo, R. Trimble, and H. Y. Mak. 2010. ’Genetic and dietary regulation of lipid droplet expansion in Caenorhabditis elegans’, Proc Natl Acad Sci U S A, 107: 4640–5.

